# Male White-shouldered Fairywrens (*Malurus alboscapulatus*) elevate testosterone when courting females but not during territorial challenges

**DOI:** 10.1101/2021.10.06.463071

**Authors:** Jordan Boersma, John Anthony Jones, Erik D. Enbody, Joseph F. Welklin, Serena Ketaloya, Jordan Karubian, Hubert Schwabl

**Affiliations:** School of Biological Sciences, Washington State University, Pullman, WA, USA; Department of Ecology and Evolutionary Biology, Tulane University, New Orleans, LA, USA; Department of Medical Biochemistry and Microbiology, Uppsala University, Uppsala, Sweden; Department of Neurobiology and Behavior, Cornell University, USA; Cornell Lab of Ornithology, Ithaca, NY, USA; Porotona Village, Milne Bay Province, Papua New Guinea

## Abstract

Testosterone mediates suites of physical and behavioral traits across vertebrates, and circulation varies considerably across and within taxa. However, an understanding of the causal factors of variation in circulating testosterone has proven difficult despite decades of research. According to the challenge hypothesis, agonistic interactions between males immediately prior to the breeding season produce the highest levels of testosterone measured during this period. While many studies have provided support for this hypothesis, most species do not respond to male-male competition by elevating testosterone. As a result, a recent revision of the hypothesis (‘challenge hypothesis 2.0’) places male-female interactions as the primary cause of rapid elevations in testosterone circulation in male vertebrates. Here, we offer a test of both iterations of the challenge hypothesis in a tropical bird species. We first illustrate that male White-shouldered Fairywrens (*Malurus alboscapulatus*) differ by subspecies in plasma testosterone concentrations. Then we use a social network approach to find that males of the subspecies with higher testosterone are characterized by greater social interaction scores, including more time aggregating to perform sexual displays. Next, we use a controlled experiment to test whether males respond to simulated territorial intrusion or courtship interaction contexts by elevating testosterone. Males sampled during courtship had greater plasma testosterone both relative to flushed controls and males sampled during simulated intrusion. Ultimately, our results are consistent with challenge hypothesis 2.0, as males rapidly elevated testosterone following interactions with females, but not during territorial challenges.

## 1. Introduction

Hormones serve a fundamental role in translating genotype to phenotype, and thus influence a population’s response to selection (Cox et al., 2016). Circulating testosterone often mediates expression of alternate male phenotypes (Ketterson and Nolan Jr, 1999; McGlothlin and Ketterson, 2008; Williams, 2012). Specifically, testosterone is responsible for many of the most conspicuous physical and behavioral traits in nature, from gaudy ornaments (Johns, 1964; Karubian et al., 2011; Peters et al., 2000; van Oordt and Junge, 1934) to aggression (reviewed in Soma 2006) and sexual display behaviors (Day et al., 2007, 2006; Fusani, 2008; Fusani et al., 2007). For these reasons, testosterone has been referred to as a phenotypic integrator that regulates a network of traits (reviewed in Lipshutz et al. 2019). While consequences of elevated testosterone circulation are reasonably well understood thanks to testosterone-implant studies (Boersma et al., 2020; Gerlach and Ketterson, 2013; Lindsay et al., 2011; McGlothlin et al., 2004; Peters, 2007; Peters et al., 2000; Peterson et al., 2013; Veiga et al., 2004; Zysling et al., 2006), the causes that lead to elevated endogenous levels in natural systems are more difficult to disentangle.

The social environment is a major source of variation in testosterone circulation across male vertebrates (reviewed in Oliveira 2004; Goymann 2009) and seasonal changes in the nature and intensity of social interactions are often attributed to temporal patterns, such as seasonal variation, of testosterone circulation across vertebrates (see Goymann et al. 2007; Goymann 2009). Wingfield et al. (1990) originally proposed the challenge hypothesis to explain the seasonal elevation of testosterone levels from a low, non-breeding baseline to the highest levels immediately prior to the breeding season. In this first iteration of the challenge hypothesis, elevated testosterone was attributed to the aggressive male-male interactions that occur during this period as males compete for breeding territories (Goymann, 2009; Wingfield et al., 1990). Whereas there is abundant empirical support for testosterone mediating aggression in male vertebrates (reviewed in Soma 2006), most species do not respond to territorial challenges by increasing circulating testosterone (Goymann et al., 2007). In contrast, interactions with fertile females are often associated with elevated testosterone in males, which provides the foundation for the revised ‘challenge hypothesis 2.0’ (Goymann et al., 2019). According to this hypothesis, interactions with receptive females are more important than intrasexual aggressive interactions for seasonal elevation of circulating testosterone in males. Empirical tests of both iterations of the challenge hypothesis within a single taxon are lacking and can help resolve the social contexts underlying rapid elevations of testosterone in males.

Moreover, our understanding of the causal factors underlying variation in testosterone is currently limited by a historical bias toward temperate systems. Tropical species tend to exhibit lower peak testosterone circulation than temperate counterparts (reviewed in Goymann et al., 2004), though there is considerable variation in testosterone in tropical birds (Moore et al., 2019). Among tropical birds greater peak testosterone is associated with higher altitudes, seasonal territory defense and breeding, as well as monogamous mating systems (Goymann et al., 2004; Hau et al., 2008). How social challenges affect circulating testosterone in tropical species is poorly understood and resolving this is necessary to improve the generalizability of hypotheses about causes of male testosterone variation across taxa.

Here we use a combination of long-term correlative studies and a controlled experiment to test both iterations of the challenge hypothesis in males of a tropical species. White-shouldered fairywrens (*Malurus alboscapulatus*) are split into six subspecies on the basis of female ornamentation, including *Malurus alboscapulatus moretoni* (hereafter: *moretoni*) which has ornamented females resembling males and *Malurus alboscapulatus lorentzi* (hereafter: *lorentzi*) which has unornamented females (Enbody et al., 2019). Unlike females, males of these subspecies are similarly ornamented across their range with no discernable variation in plumage coloration (Enbody et al. 2019). Previous studies have demonstrated that plasma testosterone is higher in *moretoni* females and that both sexes of *moretoni* are more territorial during simulated territorial intrusions (STIs) than *lorentzi* (Enbody et al., 2018). Male testosterone levels in these two subspecies are higher than in females, but although more variable than females, differences among subspecies in males were not found to be statistically different (Enbody et al., 2018). Both subspecies of White-shouldered Fairywrens defend territories and nest year-round, and are socially monogamous with biparental care of offspring, but differ in density and group size, suggesting differences in sociality (Enbody et al. 2019). Here we expand on initial comparative work to compare males of the *moretoni* and *lorentzi* subspecies in circulating testosterone and several metrics of social interaction. Then we use an experiment to test which social interaction contexts, specifically agonistic territorial intrusions or courtship opportunities, cause elevated testosterone in males.

## 2. Methods

### 2.1 Study system

White-shouldered fairywrens (Maluridae) are savanna and second-growth specialists endemic to New Guinea (Schodde, 1982). White-shouldered Fairywrens defend territories and can nest year-round, with females building nests and incubating eggs and males assisting with offspring provisioning (Enbody et al. 2019). We studied three populations of the *moretoni* subspecies, where females are ornamented like males (monochromatic plumage), and two populations of the *lorentzi* subspecies, where females are unornamented (dichromatic plumage). Our work with *moretoni* was conducted around Garuahi (150°29′ E, 10°13′ S, 0–10 m a.s.l.), Podagha (149°90′ E, 9°69’ S, 50-60 m) and Porotona, Milne Bay Province (150° 35′ E, 10°15′ S, 10–20 m a.s.l.). We conducted work with *lorentzi* near Obo village, Western Province: one population was along the western edge of the Fly River drainage (141°19′ E, 7°35′ S, 10–20 m a.s.l.), and the other was on a chain of islands in the middle of the Fly River (141°16′ E, 7°34′ S, 10–20 m a.s.l.). We used mistnets to catch Fairywrens and used a unique Australian Bird and Bat Banding Scheme metal band in addition to a unique combination of three plastic color bands to differentiate individuals for behavioral observations. Capture and behavioral work was approved by the Institutional Animal Care and Use Committee (IACUC) at Washington State University (protocol #: ASAF-04573).

### 2.2 Long-term testosterone and sociality study

#### Testosterone sampling

We regularly caught male White-shouldered Fairywrens and took blood samples from the jugular vein within 10 minutes of capture to measure plasma testosterone concentrations. Long-term work spanned field seasons from 2014-2019 across most months of the year and between the hours of 5am-noon and 3pm to 7pm when Fairywren activity was highest. Individuals that were exposed to playback prior to capture or any other experimental work were excluded from analysis. In addition, we excluded any males who were lacking cloacal protuberances (indicative of a non-breeding state) from our dataset so that we could compare males in breeding condition across subspecies.

#### Social network observations

We conducted social network observations by following a focal group for a minimum of 10 and maximum of 25 minutes during field seasons in 2016 and 2017. Observations began when we first located the focal group, and focal groups were selected each day based on the duration of time since the last observation. Focal groups were generally mated pairs, and in some cases included adult auxiliaries or fledged young. The number of stable groups varied across field seasons and populations included between 10-30 groups. Once we found the focal group we recorded spatial location data (via GPS) and the color bands for each individual in the group, repeating every 5 minutes. Individuals within 5m of each other were counted as associating. We took note of whether males aggregated and performed sexual displays to females. We identified three distinct types of male sexual displays in this species: ‘petal-carrying’ where a male carries an orange or red flower petal, leaf, or berry; ‘puff-shoulder’ where males erect shoulder patches and hop between perches near a female; and ‘display flights’ where males erect shoulder patches and back feathers and perform a bobbing flight away from the target female’s perch. If these behaviors were observed among multiple males with a female present each male in the group was counted as attending a display flock. These display aggregations resemble a lek, though there is no designated display area as in traditional lek breeding systems (Boersma, pers. obs.).

### 2.3 Sampling context experiment

#### Overall design

To test among challenge hypothesis 1.0 and 2.0 we sampled males during a simulated territorial intrusion (STI) or while they were actively courting a female. We captured a subset of males by flushing into mistnets apart from any social stimulus to serve as a control. Individual males from all populations were randomly selected to be sampled either during an STI or flushed without a stimulus. Additionally, males of the *lorentzi* subspecies were captured following female only playback and mount phenotype corresponding to the local subspecies to draw males into a courtship competition. We only sampled *lorentzi* males in the courtship competition context due to our inability to induce display flocks in *moretoni* using female playback. In all cases individuals included in this experiment had an initial blood sample taken within 10 minutes of capture (mean = 4.62 min, range = 2-10 min). In total we captured 17 *moretoni* males and 8 *lorentzi* males by flushing, 14 *moretoni* males and 13 *lorentzi* males following an STI, and 16 *lorentzi* males during a courtship competition.

#### Flush capture

Flush captures were used to assess baseline (presumably not influenced by our activities) testosterone levels. During flush captures no playback was used and we avoided rapid movements toward Fairywrens that may elicit a stress response. We ensured that no conspecifics were on a focal pair’s territory prior to capture; any instances of intrusions by conspecifics led to samples being excluded.

#### Simulated territorial intrusion (STI) capture

We used an established simulated territorial intrusion protocol for our species (for full details see Enbody et al. 2018) to sample testosterone during an agonistic challenge. Briefly, we randomly selected a cardstock White-shouldered fairywren mount pair (n = 4 exemplars of each sex) that matched the local phenotypes and played recorded duets (n = 8 exemplars) from the local subspecies. Duet playbacks were previously recorded using a protocol outlined in Enbody et al. (2018). After 10 minutes of playback we opened a furled mistnet that was placed approx. 5 m from the mounts to capture males. We recorded the time between playback start and capture to assess the effect of playback time on testosterone levels (mean playback time: 20.9 min.; range: 11-77 min.).

#### Courtship competition capture

A female mount (N = 4 separate mounts randomly assigned) painted with the local phenotype was placed on a known territory and a randomly selected female solo song recorded from the same subspecies (N = 4 individual recordings) was used to lure males from neighboring territories into forming a display flock (described in ‘social network observations’ section). Mistnets were setup near the mount, and we confirmed that males were either actively displaying or were bringing food (generally small invertebrates) to either the mount or the territorial female immediately prior to capture.

### 2.4 Testosterone radioimmunoassay methods

At the end of each field day, we centrifuged blood samples to separate plasma from red blood cells and used a Hamilton^™^ syringe to transfer plasma into 0.4 ml of 200 proof Ethanol (Fisher Bioreagants^™^) in individual Eppendorf^™^ tubes. Samples were stored at room temperature in the field, then transferred to a refrigerator in our lab at Washington State University. Prior to assay we sorted samples randomly across assays to account for inter-assay variation. We used an established radioimmunoassay protocol that measures total androgen titres instead of testosterone specifically (full details in Lindsay et al., 2011) because the antibody we use (Wien Laboratories T-3003, Flanders, NJ, USA) cross-reacts strongly with 5α-dihydrotestosterone (DHT). Long-term testosterone dataset samples were split among 15 separate assays with an inter-assay coefficient of variation of 17.29% (calculated following methods in Chard 1995). The experimental sampling context samples were split among 5 separate assays with an inter-assay of 13.57%. Plasma volumes ranged from 19.94 – 66.23 µl (mean = 39.51 µl) in our long-term sample pool, and 12.14 – 28.37 µl (mean = 22.43 µl) in our experimental sample pool. The minimum detectable androgen concentration calculated based on a 12 µl (lowest) plasma volume and mean recovery rate of 65.77% across assays was 308.86 pg/ ml. We used our assay’s minimal detectable level of 1.95 pg/ tube to assign a total androgen concentration among undetectable samples (N = 23 in long-term dataset, N = 7 in experimental dataset).

### 2.5 Statistical methods

All analyses were conducted in R (<www.r-project.org>) version 3.6.1 using linear regressions in base R and mixed models in package lme4 (Bates et al., 2015). We used backward stepwise selection following Wang et al. (2008) throughout: the full set of candidate predictors was initially included in each model and any variable with p ≥ 0.4 was removed. Post-hoc Tukey tests were used following detection of a significant effect when there were more than two treatment levels using package multcomp (Hothorn et al., 2008). We used a permutation-based approach for both long-term testosterone and social network analyses following Farine (2017). In short, we compared the observed model coefficients for the subspecies variable to the distribution of coefficients from 1,000 randomized datasets where subspecies identity had been randomly reassigned. When randomizing subspecies identity, we required that individuals sampled multiple times were assigned the same subspecies identity in each of the randomized datasets. *Long-term testosterone*

To improve normality for statistical analysis, we natural log (ln) transformed circulating testosterone concentrations (pg/ ml plasma) and inspected residuals with normal Q-Q plots. We used a linear mixed model with ln testosterone as the response variable and subspecies, year, time of day bled, net-capture-to-bleed time, and nesting status (confirmed active nest or unknown) and date as fixed effects in the initial model. To account for repeated measures, we included individual ID as a random effect.

#### Social network analysis

We used R package asnipe (Farine, 2013) to build social networks using the simple ratio index (SRI). We used the igraph package (Csardi and Nepusz, 2006) to detect and plot social network structure and the program Gephi (Bastian et al., 2009) to depict where individuals were observed using GPS data associated with each observation. We compiled data across both sampling years to generate the social network plot, but split data for each male by sampling year in our models. There were 1,005 separate observations made in *lorentzi* (681 in 2016 and 324 in 2017) and 795 observations in moretoni (465 in 2016 and 330 in 2017). Prior to running models we removed individuals from the dataset that were observed less than 5 times in a sampling year. This resulted in 47 total *lorentzi* males and 43 *moretoni* males observed across years, with 28 *lorentzi* males and 23 *moretoni* males observed in 2016 and 19 *lorentzi* males and 20 *moretoni* males observed in 2017.

To determine how subspecies differ in sociality, we calculated five social metrics for each male: (1) unweighted male degree, or the number of conspecific males an individual interacted with; (2) unweighted female degree, or the number of conspecific females an individual interacted with; (3) weighted male degree, or the number of male conspecifics interacted with weighted by the total number of interactions; (4) weighted female degree, or the number of female conspecifics interacted with weighted by the total number of interactions; and (5) the number of display flocks each individual male took part in during the sampling period (full details in “social network observations” section). We chose to analyze both unweighted and weighted degree scores because they capture different aspects of social behavior: unweighted degree indicates how many social partners each male has, while weighted degree reflects the strength of those partnerships. Each metric was analyzed in separate linear models with subspecies, year, and sight frequency (the # of times an individual was observed) as fixed effects. Individual ID was included as a random effect to account for repeated measures.

#### Testosterone across experimental sampling contexts

As above, we used a natural log (ln) transformation for testosterone and checked for normality of residuals. We used two separate linear mixed models with testosterone as the response variable. Both had experimental sampling context (flush, STI, courtship), subspecies, time of day bled, net-to-bleed time, and nesting status (nesting, not nesting, or unknown based on state of female’s brood patch) as fixed effects with individual ID as a random effect. We first checked for an interaction between subspecies and experiment with the courtship competition samples excluded because we lacked trials in *moretoni* subspecies. Then we used a separate model to compare all levels of our sampling context variable with the non-significant interaction excluded (interaction model terms in Table S2).

## 3. Results

### 3.1 Long-term study

#### Testosterone

We sampled plasma testosterone in 42 individual *lorentzi* and 58 *moretoni* males for a total of 68 and 92 samples, respectively. Log transformed testosterone significantly differed according to subspecies, breeding status, and bleed time (Figure 1; full model terms: Table 1). The *lorentzi* subspecies had higher plasma testosterone than the *moretoni* subspecies (*p* = 0.008, *lorentzi* mean: 1118.75 pg/ ml; *moretoni* mean: 660.19 pg/ ml). Males known to have an active nest (N = 19) had higher testosterone than males without a known nest (*p* = 0.028; active nest mean: 1320.73 pg/ ml; unknown nest status mean: 792.33 pg/ ml). There was a positive relationship between circulating testosterone and time of day bled (*p* = 0.024). Net-to-bleed time and year had no effect (*p* > 0.4) and removed from the model.

**Table 1.** Model results for linear mixed model analysis of male plasma testosterone from our long-term dataset. Bolded values show significant effects and italics depict terms removed from the final model.

| Long-term log transformed testosterone (N = 160 samples, 100 males) |  |  |  |  |  |  |
| --- | --- | --- | --- | --- | --- | --- |
| Predictor | $\beta$ | SE | $t$ | df | $p$ | Random effects $\sigma$ |
| Intercept | 5.772 | 0.317 | 18.204 |  |  | Male ID <0.001 |
| Subspecies | -0.405 | 0.173 | -2.355 | 1 | **0.008** |  |
| Nesting status | 0.577 | 0.266 | 2.171 | 1 | **0.030** |  |
| Time of day bled | 0.044 | 0.020 | 2.226 | 1 | **0.026** |  |
| *Net-to-bleed time* | *0.009* | *0.043* | *0.199* | *1* | *0.842* |  |
| *Date* | *<0.001* | *<0.001* | *-0.283* | *1* | *0.777* |  |

**Figure 1.**
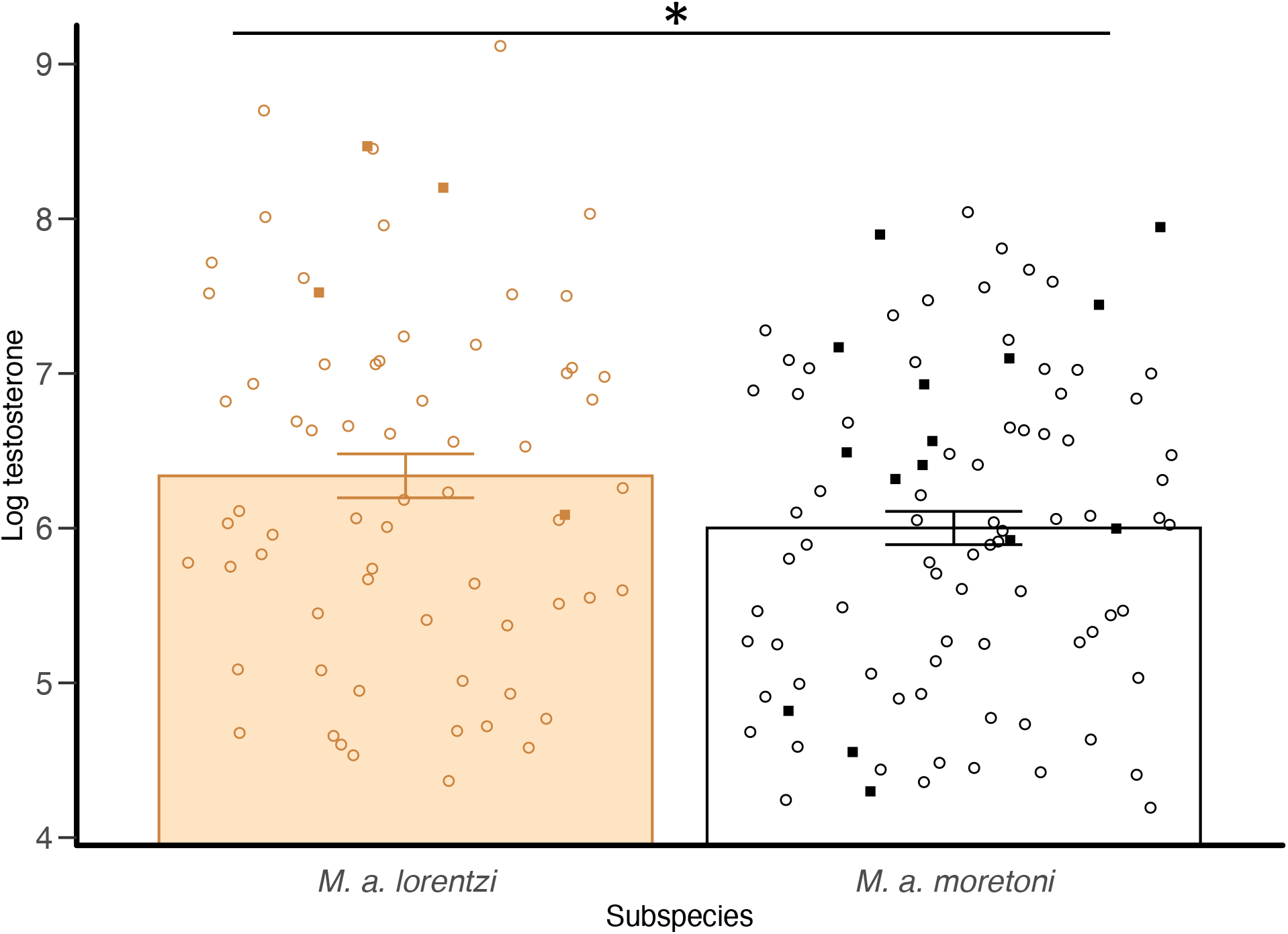
Testosterone by subspecies. Natural log-transformed testosterone in males across two subspecies of White-shouldered Fairywren. Bars represent means with standard error overlaid and asterisks indicate significant difference between subspecies (* p < 0.05). All males were in breeding condition and those with active nests are depicted with filled squares.

#### Sociality

Social networks for each sampling site (subspecies) are depicted in Figure 2. Males differed significantly by subspecies in each social metric except female weighted degree (Figure 3; full model terms: Table 2). Males of the *lorentzi* subspecies were characterized by higher female degree (perm *p* <0.001; *lorentzi* mean: 2.26, *moretoni* mean: 1.02) and male degree scores (perm *p* <0.001; *lorentzi* mean: 2.51, *moretoni* mean: 0.42), higher male weighted degree scores (perm *p* <0.001; *lorentzi* mean: 0.29, *moretoni* mean: 0.06), and greater display flock attendance (perm *p* <0.001; *lorentzi* mean: 0.85, *moretoni* mean: 0.12) than the *moretoni* subspecies, but no difference in female weighted degree (perm *p* = 0.53; *lorentzi* mean: 0.63, *moretoni* mean: 0.63). Sight frequency was not a significant predictor in any model.

**Table 2.** Model results for linear model analysis of each social metric across male subspecies. Bolded values depict significant differences, and italicized

| **(a) Female degree (N = 51 males, 90 observations)** |  |  |  |  |  |  |  |
| --- | --- | --- | --- | --- | --- | --- | --- |
| Predictor | $\beta$ | SE | *t* | *df* | *p* | Random effects | $\sigma$ |
| Intercept | 1.644 | 0.310 | 5.305 |  |  | Male ID | <0.001 |
| Subspecies | -1.122 | 0.265 | -4.240 | 1 | **<0.001** |  |  |
| Sight frequency | 0.0354 | 0.015 | 2.426 | 1 | **0.015** |  |  |
| Year | -0.105 | 0.278 | -0.377 | 1 | 0.706 |  |  |
| **(b) Male degree (N = 51 males, 90 observations)** |  |  |  |  |  |  |  |
| Predictor | $\beta$ | SE | *t* | *df* | *p* | Random effects | $\sigma$ |
| Intercept | 2.511 | 0.211 | 11.896 |  |  | Male ID | <0.001 |
| Subspecies | -2.092 | 0.305 | -6.852 | 1 | **<0.001** |  |  |
| Sight frequency | -0.006 | 0.018 | -0.357 | 1 | 0.721 |  |  |
| Year | 0.234 | 0.325 | 0.720 | 1 | 0.471 |  |  |
| **(c) Female weight. degree (N = 51 males, 90 observations)** |  |  |  |  |  |  |  |
| Predictor | $\beta$ | SE | *t* | *df* | *p* | Random effects | $\sigma$ |
| Intercept | 0.625 | 0.0629 | 9.943 |  |  | Male ID | 0.152 |
| Subspecies | 0.005 | 0.090 | 0.058 | 1 | 0.530 |  |  |
| Sight frequency | 0.002 | 0.005 | 0.407 | 1 | 0.684 |  |  |
| Year | 0.065 | 0.089 | 0.727 | 1 | 0.467 |  |  |
| **(d) Male weighted degree (N = 51 males, 90 observations)** |  |  |  |  |  |  |  |
| Predictor | $\beta$ | SE | *t* | *df* | *p* | Random effects | $\sigma$ |
| Intercept | 0.311 | 0.066 | 4.691 |  |  | Male ID | 0.084 |
| Subspecies | -0.249 | 0.050 | -4.981 | 1 | **<0.001** |  |  |
| Sight frequency | -0.004 | 0.003 | -1.332 | 1 | 0.183 |  |  |
| Year | 0.111 | 0.0478 | 2.331 | 1 | **0.020** |  |  |
| **(e) Display flock attend. (N = 51 males, 90 observations)** |  |  |  |  |  |  |  |
| Predictor | $\beta$ | SE | *t* | *df* | *p* | Random effects | $\sigma$ |
| Intercept | 0.850 | 0.100 | 8.525 |  |  | Male ID | 0.107 |
| Subspecies | -0.733 | 0.144 | -5.085 | 1 | **<0.001** |  |  |
| Sight frequency | 0.002 | 0.008 | 0.275 | 1 | 0.783 |  |  |
| Year | -0.055 | 0.152 | -0.36 | 1 | 0.719 |  |  |

**Figure 2.**
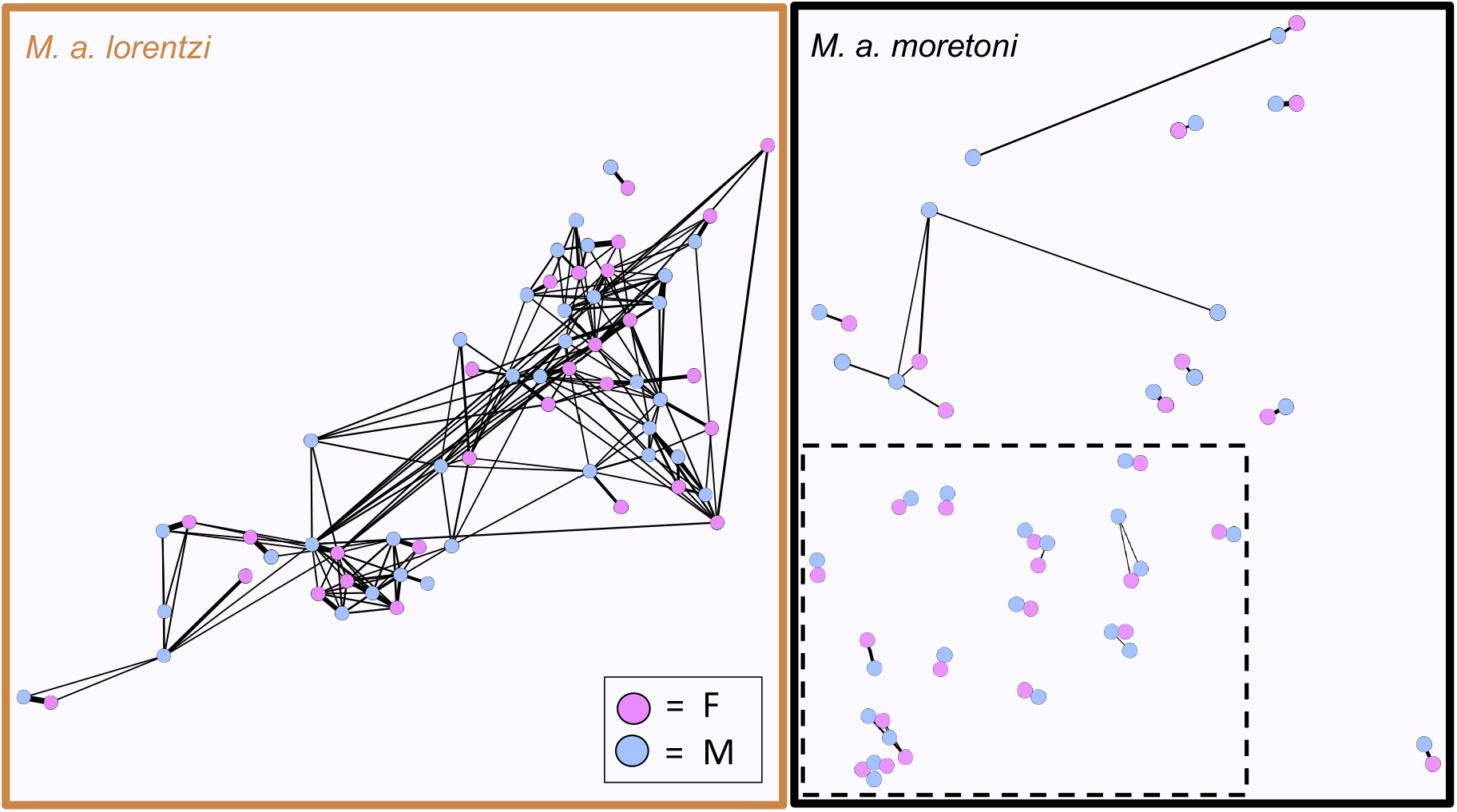
Social networks by subspecies. Social network structure across two subspecies of White-shouldered Fairywren. Dashed inset (bottom right) separates the two populations of *M. a. moretoni* used for comparisons. Each node is an adult male (blue circle) or female (pink) observed 10 or more times over the study period (*lorentzi*: 26 males and 21 females; *moretoni*: 25 males and 21 females). Lines between individual represent interactions, and thickness of each line depicts the frequency in which individuals interacted.

**Figure 3.**
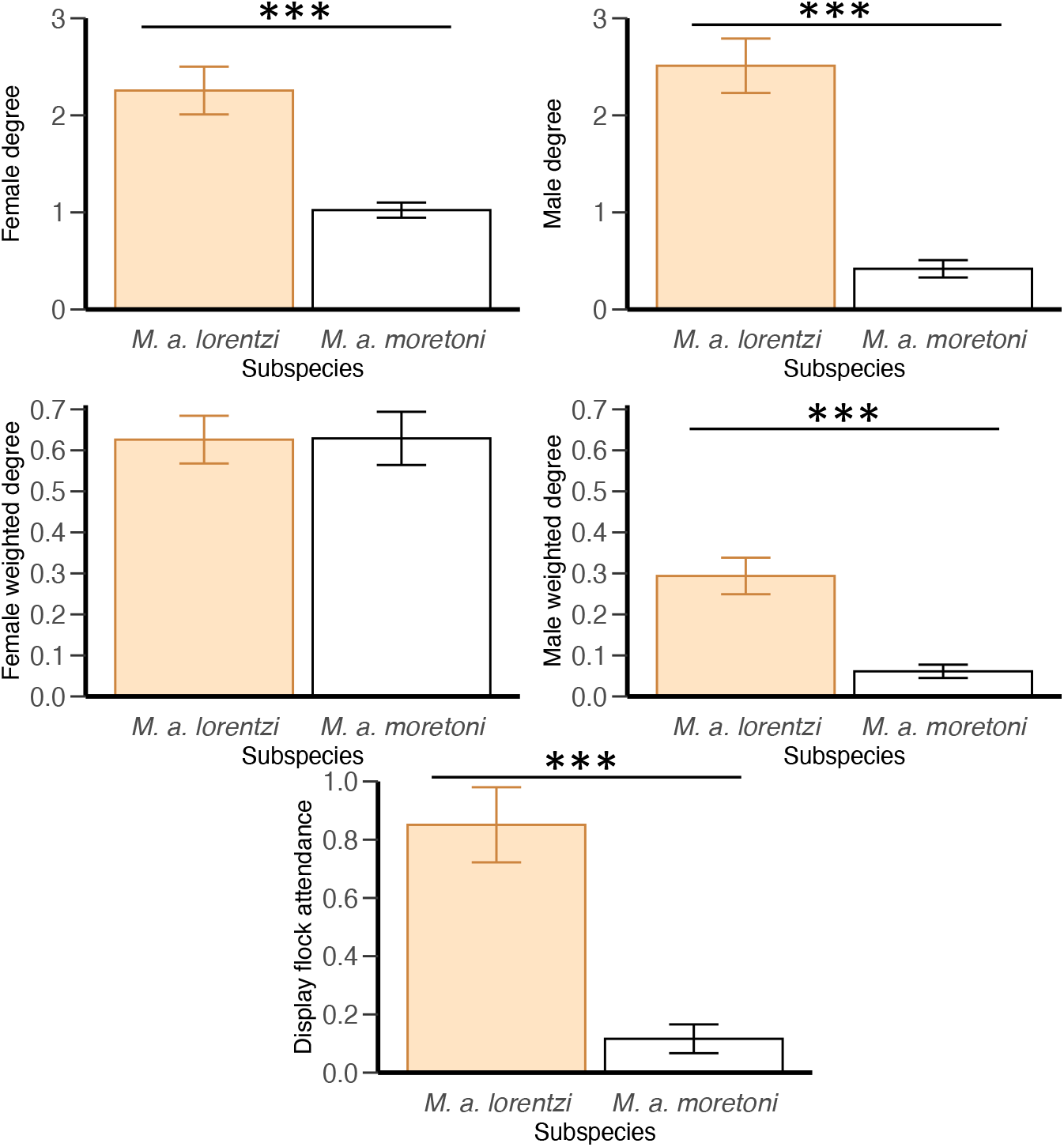
Social interaction by subspecies. Social interaction score (across subspecies. Bars represent means with standard error overlaid and asterisks indicate significant comparisons (***p < 0.001).

### 3.2 Testosterone across experimental sampling contexts

To evaluate if males of the two subspecies responded differently to sampling context, we first tested whether there was an interaction between subspecies and sampling context (flush and STI) on plasma testosterone and did not find a significant effect (*p* = 0.57; Table S2). We then analyzed whether plasma testosterone differed by sampling context with all three levels included (flush, STI, courtship competition) with the interaction between subspecies and sampling context excluded. The final model included both sampling context and breeding status (pooling across subspecies), with sampling context as the only significant predictor (*p* < 0.001; full model terms: Table 3). A post-hoc Tukey test revealed that *lorentzi* males sampled in the courtship context had higher testosterone than males who were flushed into the net (Figure 4; *p* < 0.001; courtship competition mean: 1219.59 pg/ ml, flushed mean: 381.66 pg/ ml) or sampled after an STI (*p* < 0.001; mean: 601.86 pg/ ml). Males sampled during an STI did not differ significantly from males who were flushed into the net without playback (*p* = 0.10). Duration of exposure to playback was not predictive of testosterone following an STI (*p* = 0.83; Table S3).

**Table 3.** Model results for linear mixed model analysis of testosterone across experimental sampling contexts. Bolded values show significant effects and italics depict terms removed from the final model. Values for experiment are in reference to the mating context and nesting status is in reference to males paired to females without active brood patches.

| **Experimental log transformed testosterone (N = 68 samples, 63 males)** |  |  |  |  |  |  |
| --- | --- | --- | --- | --- | --- | --- |
| **Predictor** | **$\beta$** | ***SE*** | ***t*** | ***df*** | ***p*** | **Random effects $\sigma$** |
| Intercept | 7.475 | 0.244 | 30.587 |  |  | Male ID <0.001 |
| Experiment (Flush) | -1.398 | 0.244 | -5.475 | 2 | **<0.001** |  |
| Experiment (STI) | -0.994 | 0.255 | -3.884 | 2 | **<0.001** |  |
| *Subspecies* | *0.151* | *0.227* | *0.667* | *1* | *0.505* |  |
| Nesting status (unknown) | -0.208 | 0.244 | -0.852 | 2 | 0.210 |  |
| Nesting status (yes) | 0.292 | 0.238 | 1.228 | 2 | 0.210 |  |
| *Time of day bled* | *-0.003* | *0.030* | *-0.089* | *1* | *0.929* |  |
| *Net-to-bleed time* | *<0.001* | *0.043* | *0.008* | *1* | *0.993* |  |
| **Post-hoc comparisons** |  |  |  |  |  |  |
| Experiment | **$\beta$** | ***SE*** | ***z*** | ***p*** |  |  |
| Flush vs. Courtship | -1.398 | 0.255 | -5.475 | **<0.001** |  |  |
| STI vs. Courtship | -0.994 | 0.256 | -3.884 | **<0.001** |  |  |
| STI vs. Flush | 0.404 | 0.198 | 2.037 | 0.102 |  |  |

**Figure 4.**
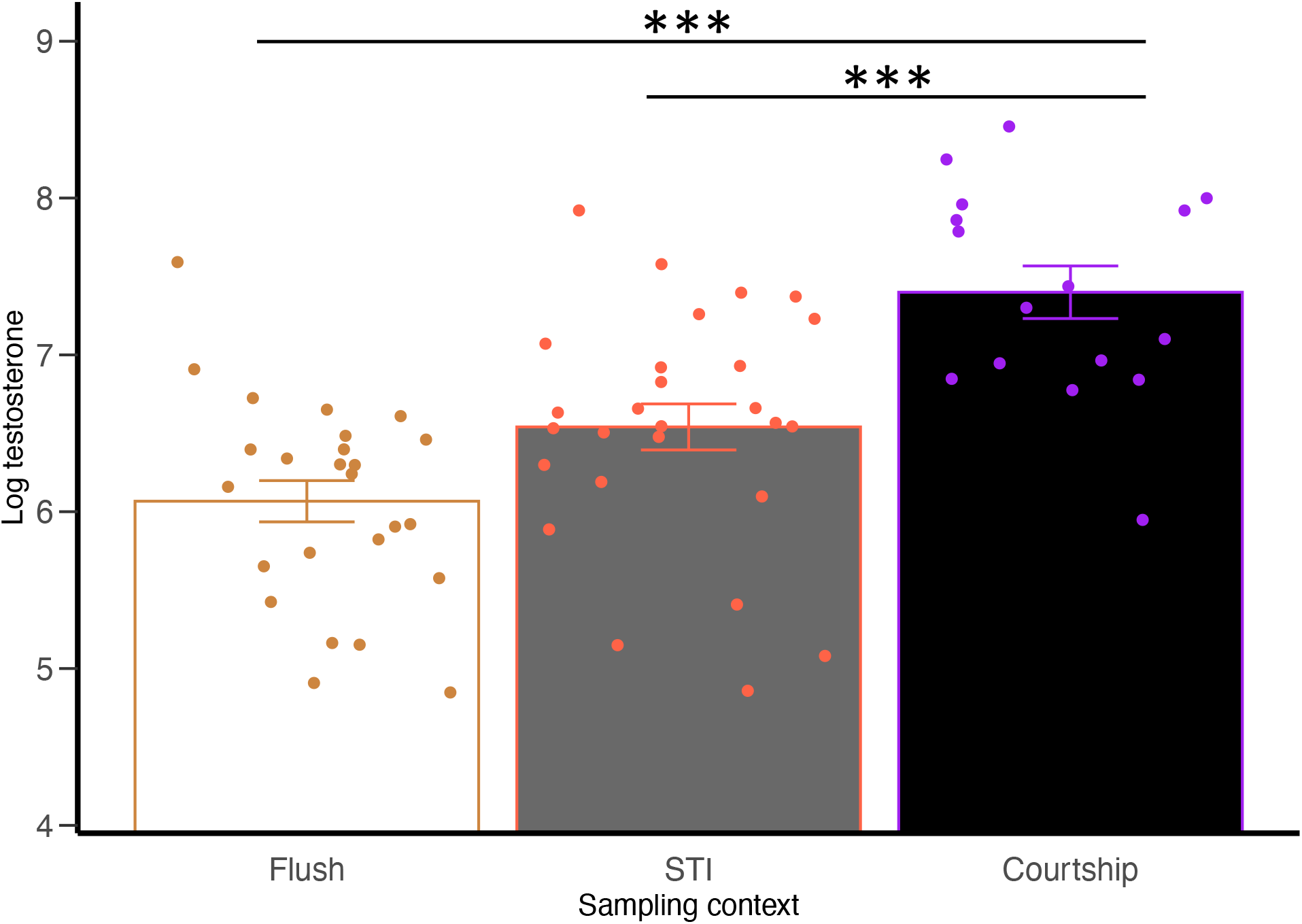
Testosterone across experimental sampling contexts. Natural log-transformed testosterone across experimental sampling context. Bars represent means with standard error overlaid and asterisks indicate significant comparisons (*** p < 0.001).

## 4. Discussion

We used a combination of a long-term correlative study and controlled experiments to test which factors are associated with elevated circulating testosterone in male White-shouldered Fairywrens of different subspecies. First, we found that males of the *lorentzi* subspecies (males ornamented, females un-ornamented) have higher circulating testosterone and are also more social than *moretoni* males (males and females ornamented) in four out of five metrics of sociality. We then tested how different social interaction contexts (simulated territorial intrusion (STI) and courtship competition) affect testosterone relative to flushed controls. We found that males sampled while actively courting females had significantly greater circulating testosterone than flushed control males or those sampled during an STI in *lorentzi*. We did not find support for an effect of simulated territorial intrusions on testosterone. Ultimately, our results are consistent with the second iteration of the challenge hypothesis (Goymann et al., 2019) over the initial version (Wingfield et al., 1990), suggesting that elevated circulating testosterone is driven by active courtship of females.

Decades of research have established a prominent role for the social environment in causing variation in male testosterone circulation. However, most of these studies have focused on gross temporal/seasonal patterns of testosterone circulation in temperate organisms. Our correlative study adds to a growing body of literature illustrating a link between both social organization and finer-grained social interactions and testosterone (Gleason et al., 2009; Goymann et al., 2019, 2004; Maguire et al., 2021; Oliveira, 2004; Ryder et al., 2011). Males of the *lorentzi* subspecies exhibited greater mean testosterone circulation than *moretoni* males across six study years (Figure 1). Although the previous study of male testosterone in this species did not find differences by subspecies (Enbody et al., 2018), our sample size is much larger for *lorentzi* (N = 26 in previous study vs. N = 68 in current study), and here we exclude the potential confound of playback use prior to capture. Interestingly, female White-shouldered Fairywrens showed the opposite pattern by subspecies: *lorentzi* females had lower mean plasma testosterone than *moretoni* females in the previous study of testosterone in this species (Enbody et al., 2018). However, this has been attributed to variation in female ornamentation, as *moretoni* females exhibit male-like ornamentation, while *lorentzi* do not, although, they do produce some ornamentation when given exogenous testosterone (Boersma et al., 2020). In the current study *lorentzi* males were found to have higher circulating testosterone levels and greater social interaction scores, including interaction with more individual males and females and aggregating in display flocks (Figures 2 and 3), which is consistent with social interactions driving subspecific male testosterone levels. Due to the transient nature of testosterone elevation and that our social data are aggregated across days without testosterone sampling, we do not present a within-population correlation between testosterone and sociality here.

We used an experiment to test which social interaction contexts affect testosterone circulation. Males sampled while actively courting a female had significantly greater testosterone circulation than those sampled while defending a territory from a simulated intruder or after being flushed into the net without a social stimulus (Figure 4). This is the first direct test to our knowledge of challenge hypothesis 1.0 vs. 2.0 (Goymann et al., 2019; Wingfield et al., 1990), and our results are consistent with the more recent iteration. Previous studies have illustrated that testosterone is elevated in males during courtship, thus informing the amended, “female interaction focused”, 2.0 version of the challenge hypothesis (Goymann et al., 2019). Our study is another in a myriad of studies that fail to find support for the original, “agonistic focused”, challenge hypothesis (meta-analysis in Hirschenhauser and Oliveira, 2006). Nesting status was not predictive of a male’s testosterone response across experimental sampling context, suggesting that the social context a male is sampled in is more important than whether he is actively providing paternal care or defending a mate. However, without details on the specific nesting stage we cannot exclude the possibility that parental versus sexual phases of nesting contributed to variation in testosterone in our dataset. It is also possible that we failed to elicit a significant hormonal response to simulated territorial intrusion because intrusions were not of long enough duration, as males of some tropical species may elevate testosterone only after 2 hours (Wikelski et al., 1999). The longest intrusion test in our study lasted 77 minutes, with 4 males sampled after at least 30 minutes of playback, but there was no relationship between playback time and testosterone following an agonistic challenge.

The drivers of variation in testosterone levels in tropical species is poorly understood (Goymann et al., 2004; Hau et al., 2008b; Moore et al., 2019). Comparative analyses of male tropical species suggest that testosterone is typically low unless species have short breeding seasons and/or experience great social instability (Goymann et al., 2004; Hau et al., 2008b; Wikelski et al., 1999). Cooperatively breeding species, which are more common in the tropics (Feeney et al., 2013), can exhibit relatively high testosterone levels due to competition among unrelated breeders and helpers for limited mating opportunities (Khan et al., 2001; Peters et al., 2001; Poiani and Fletcher, 1994; Vleck and Brown, 1999; Wingfield et al., 1991). Likewise, tropical lek breeding species where males court females throughout much of the year seem to have testosterone levels comparable to temperate species (Fusani et al., 2007; Moore et al., 2019; Ryder et al., 2011), and testosterone can be associated with and stimulate courtship display behavior (Chiver and Schlinger, 2017; Day et al., 2007, 2006; Wiley and Goldizen, 2003). Male White-shouldered Fairywrens exhibit life history characteristics consistent with both high and low testosterone tropical species. They defend territories and maintain breeding readiness year-round, with long-term pair bonds and biparental care, with some level of cooperative breeding and aggregation at mobile lek display sites (Enbody et al., 2019). Given the *moretoni* subspecies with ornamented females shows more aggressive territorial defense (Enbody et al., 2018), our finding that *lorentzi* males are characterized by more social interaction, including greater rates of sexual displays to females, indicate potentially stark differences in social environments across subspecies. The highest testosterone values in our long-term dataset come from *lorentzi* males, and our experimental work showed that *lorentzi* males sampled while actively courting females had higher testosterone than those sampled during an agonistic intrusion or absent a social stimulus. Collectively, our results indicate that courtship interactions are responsible for elevated testosterone in males of this tropical species, supporting challenge hypothesis 2.0.

## Acknowledgements

We thank the generous landowners at each of our field sites in Papua New Guinea, without which none of our studies would be possible. Many individuals assisted with field work, and we’d like to thank J. A. Gregg, I. R. Hoppe, D. Nason, M. Olesai, J. Sarusaruna, Kipling and Manu for their expert help in the field. We are grateful for funding from the Washington State University Elling Fund and American Ornithological Society (to J. B.), the Evolutionary Biology Department Student Research Grant and Gunning Fund (to J. A. J. and E. D. E.) and the Disney Conservation Fund (to J. K.) and National Science Foundation (IOS-1354133 to J. K. and IOS-1352885 to H. S.).

